# 25-Hydroxycholesterol amplifies microglial IL-1β production in an apoE isoform-dependent manner

**DOI:** 10.1101/2020.01.14.906966

**Authors:** Man Ying Wong, Michael Lewis, James J. Doherty, Yang Shi, Patrick M. Sullivan, Mingxing Qian, Douglas F. Covey, Gregory A. Petsko, David M. Holtzman, Steven M. Paul, Wenjie Luo

**Affiliations:** Appel Alzheimer’s Disease Research Institute, Feil Family Brain and Mind Research Institute, Weill Cornell Medicine, New York, USA; Sage Therapeutics, Cambridge, Massachusetts, USA; Department of Neurology, Hope Center for Neurological Disorders, Charles F. and Joanne Knight Alzheimer’s Disease Research Center, Washington University School of Medicine, St. Louis, Missouri, USA.; Department of Medicine, Duke University Medical Center, Durham Veterans Health Administration Medical Center’s Geriatric Research, Education and Clinical Center, Durham, NC, USA; Departments of Developmental Biology, Anesthesiology, Taylor Family Institute for Innovative Psychiatric Research, Washington University in St. Louis, 660 South Euclid Avenue, St. Louis, MO 63110, USA; Departments of Neurology and Psychiatry, Hope Center for Neurological Disorders, Taylor Family Institute, Washington University School of Medicine, St. Louis, Missouri, USA

## Abstract

Genome-wide association studies associated with Alzheimer’s disease (AD) have implicated pathways related to both lipid homeostasis and innate immunity in the pathophysiology of AD. However, the exact cellular and chemical mediators of neuroinflammation in AD remain poorly understood. The oxysterol 25-hydroxycholesterol (25-HC) is an important immunomodulator produced by peripheral macrophages with wide-ranging effects on cell signaling and innate immunity. Genetic variants of the enzyme responsible for 25-HC production, cholesterol 25-hydroxylase (CH25H), have been found to be associated with AD. In the present study, we found that the CH25H expression is upregulated in human AD brain tissue and in transgenic mouse brain tissue bearing amyloid-β (Aβ) plaques or tau pathology. Treatment with the toll-like receptor 4 (TLR4) agonist lipopolysaccharide (LPS) markedly upregulates CH25H expression in the mouse brain *in vivo*. LPS also stimulates CH25H expression and 25-HC secretion in cultured mouse primary microglia. We also found that LPS-induced microglial production of the pro-inflammatory cytokine IL1β is markedly potentiated by 25-HC and attenuated by genetic deletion of CH25H. Microglia expressing apolipoprotein E4 (apoE4), a genetic risk factor for AD, produce greater amounts of 25-HC than apoE3-expressing microglia following treatment with LPS. Remarkably, treatment of microglia with 25-HC results in a much greater level of IL1β secretion in LPS-activated apoE4-expressing microglia than in apoE2- or apoE3-expressing microglia. Blocking potassium efflux or inhibiting caspase-1 prevents 25-HC-potentiated IL1β release in apoE4-expressing microglia, indicating the involvement of caspase-1/NLRP3 inflammasome activity. 25-HC may function as a microglial secreted inflammatory mediator in brain, promoting IL1β-mediated neuroinflammation in an apoE isoform-dependent manner (E4≫E2/E3) and thus may be an important mediator of neuroinflammation in AD.

## Introduction

Neuroinflammation is a prominent feature of the neuropathology of Alzheimer’s disease (AD), in addition to β-amyloid (Aβ) plaques and tau-containing neurofibrillary tangles (NFT) (1). Emerging evidence indicates that neuroinflammation, mediated by activated glial cells, plays a fundamental role in the pathogenesis and neurodegeneration of AD (1). Brain inflammation either triggered by or proceeding AD pathology sustains and likely contributes to the progressive neurodegeneration that characterizes AD (2). Defining the molecular and cellular mechanisms underlying neuroinflammation as well as the chemical mediators of the inflammatory cascade are critical for understanding how neuroinflammation contributes to AD pathogenesis.

In AD, neuroinflammation increases with disease progression and is primarily driven by glial cells, especially microglia. This pathophysiological inflammatory cascade is associated with increased production of pro-inflammatory cytokines and other key inflammatory mediators (3, 4), including interleukin-1β (IL-1β), a very potent pro-inflammatory cytokine (5–8). Higher concentrations of IL-1β have been reported in cerebrospinal fluid and brain tissue of AD patients (9–11) and in microglia surrounding Aβplaques (12). Sustained elevations of IL-1β have been postulated to play a key role in AD pathogenesis (6, 12–14). Active IL-1β (17kD) is produced from an inactive 31 kDa pro-IL-1β by the inflammasome, a multicomponent protein complex consisting of NLRP3 (nucleotide-binding domain and leucine-rich repeat-containing protein 3), ASC (apoptosis-associated speck-like protein containing a CARD) and caspase-1(15). The elevations of IL-1β reported in AD brain strongly suggests activation of the inflammasome (16). Supporting this, aggregated Aβ has been shown to activate the inflammasome via a CD36/TLR4/6-dependent mechanism (17). NLRP3 deficiency reduces amyloid deposition and rescues memory deficits in the APP/PS1 model of AD (18). Understanding the cellular mechanisms responsible for IL-1β production by microglia may facilitate the development of an effective AD therapeutic that reduces IL-1β-mediated immune signaling and associated neuroinflammation.

The apolipoprotein E4 (APOE4) allele is the most common and important genetic risk factor for late onset AD (19–21). In the periphery, apoE regulates lipid metabolism (22, 23). In brain, apoE functions as an important regulator of brain amyloid (amyloid β-peptide or Aβ) deposition and clearance (apoE2>E3>E4), which most likely accounts for one of the known mechanisms as to how APOE4 increase AD risk (24). Recently, several studies have shown that APOE4 is associated with increased innate immune reactivity and enhanced cytokine secretion in primary microglia and peripheral macrophages in various animal models as well as human subjects (25–36). We have also reported a higher innate immune reactivity of apoE4-expressing microglia following LPS treatment and found that APOE4/4 genotype greatly influences tau-dependent neuroinflammation in a tau transgenic mouse model of neurodegeneration (37). Together, these data suggest that apoE4 may exert a “toxic” gain of function to promote microglia-mediated neuroinflammation and neurodegeneration in AD.

25-hydroxycholesterol (25-HC) is a potent oxysterol regulator of cholesterol biosynthesis (38–40). It is converted from cholesterol by the oxidoreductase cholesterol 25-hydroxylase (CH25H) (41, 42), an enzyme highly expressed and induced primarily in peripheral macrophages and dendritic cells in response to inflammatory stimuli like LPS and interferon (43, 44). Although CH25H deficiency does not cause defects in cholesterol homeostasis (44, 45), 25-HC appears to serve multiple functions to regulate both innate and adaptive immunity. It acts as either an anti-or pro-inflammatory regulator involved in protection from viral infection, macrophage foam cell formation, immunoglobin IgA production and cytokine production (44). To date, the function of CH25H and 25-HC in the central nervous system has not been well characterized. The association of CH25H with AD was first reported in a hippocampal microarray study of AD brain tissue (46) and further supported by an AlzGene meta-analysis for a sporadic AD population (47) and other AD patient-based independent systematic analyses (48–50). Upregulation of CH25H mRNA in affected brain regions in AD patients versus controls was first reported in a hippocampal microarray. Upregulation of CH25H expression has also been detected in brain tissue of AD transgenic mice (51–53). It remains unclear exactly how CH25H and its oxysterol product 25-HC are involved in AD.

In the present study, we investigated whether 25-HC regulates the innate immune response of microglia and whether the APOE4 allele relative to the other common APOE alleles impacts the effects of 25-HC on microglial activation. Our results demonstrate that CH25H is upregulated in AD brain tissue and AD transgenic mouse brain. We further show that 25-HC is produced by activated primary microglia and augments IL-1βproduction stimulated by the TLR4 agonist LPS. Importantly microglia expressing apoE4 produce much greater amounts of 25-HC and IL-1β in response to LPS treatment compared to apoE2-or apoE3-expressing microglia. Remarkably, 25-HC also markedly potentiates LPS-mediated IL-1β secretion by apoE4-expressing microglia. Inhibition of inflammasome activity markedly reduces augmentation of microglial IL-1β secretion by 25-HC. Our results suggest that 25-HC may function as an inflammatory mediator of the IL-1β-dependent inflammatory cascade in microglia and thus, may contribute to apoE4-dependent neuroinflammation and neurodegeneration in AD.

## Results

### CH25H is upregulated in human AD brain and AD-related transgenic mouse brain

We first examined the expression of CH25H in postmortem human AD brain tissue. Using quantitative PCR, we observed that the level of CH25H mRNA was significantly upregulated in frontal cortical tissue of AD brain (n=14) compared to age-matched (non-AD) controls (n=9, *p<0.05*) (Fig.1a, all subjects were age>80 and both genders included). The protein level of CH25H was also increased in AD brain tissue as detected by Western Blot using a CH25H antibody (Fig. 1b). The increased levels of CH25H mRNA and protein were also observed in the frontal cortex of 4-mo old APPPS1-21 mice bearing amyloid plaques (54) (Fig.1c, d, e). We further examined the expression of CH25H in PS19 mice expressing the human P301S tau mutation at 9-mo of age bearing massive tau pathology, inflammation and neurodegeneration in brain (55). Compared to their age-matched non-tg littermates, we detected an increase of CH25H mRNA in the brain of PS19 tg mice (Fig. 1f). Moreover, when we measured CH25H mRNA levels in the frontal cortex of P301S tau transgenic mice that are homozygous for human APOE2 (TE2), APOE3 (TE3), APOE4 (TE4) or with no expression of apoe (TEKO) using nanostring analysis, we found that TE4 mice, an aggressive mouse model showing the strongest brain neurodegeneration and neuroinflammation (37), express significantly higher levels of CH25H mRNA than TEKO mice, and also showed higher levels of CH25H expression than TE2 or TE3 mice although these comparisons did not reach statistical significance (Fig. 1g). Together these data suggest that CH25H expression is upregulated in human AD brain and mouse brain when there is prominent amyloid or tau pathology and neuroinflammation. Given that we measured CH25H mRNA and protein in brain tissue and not in individual microglia, these data are all the more noteworthy.

**Figure 1:**
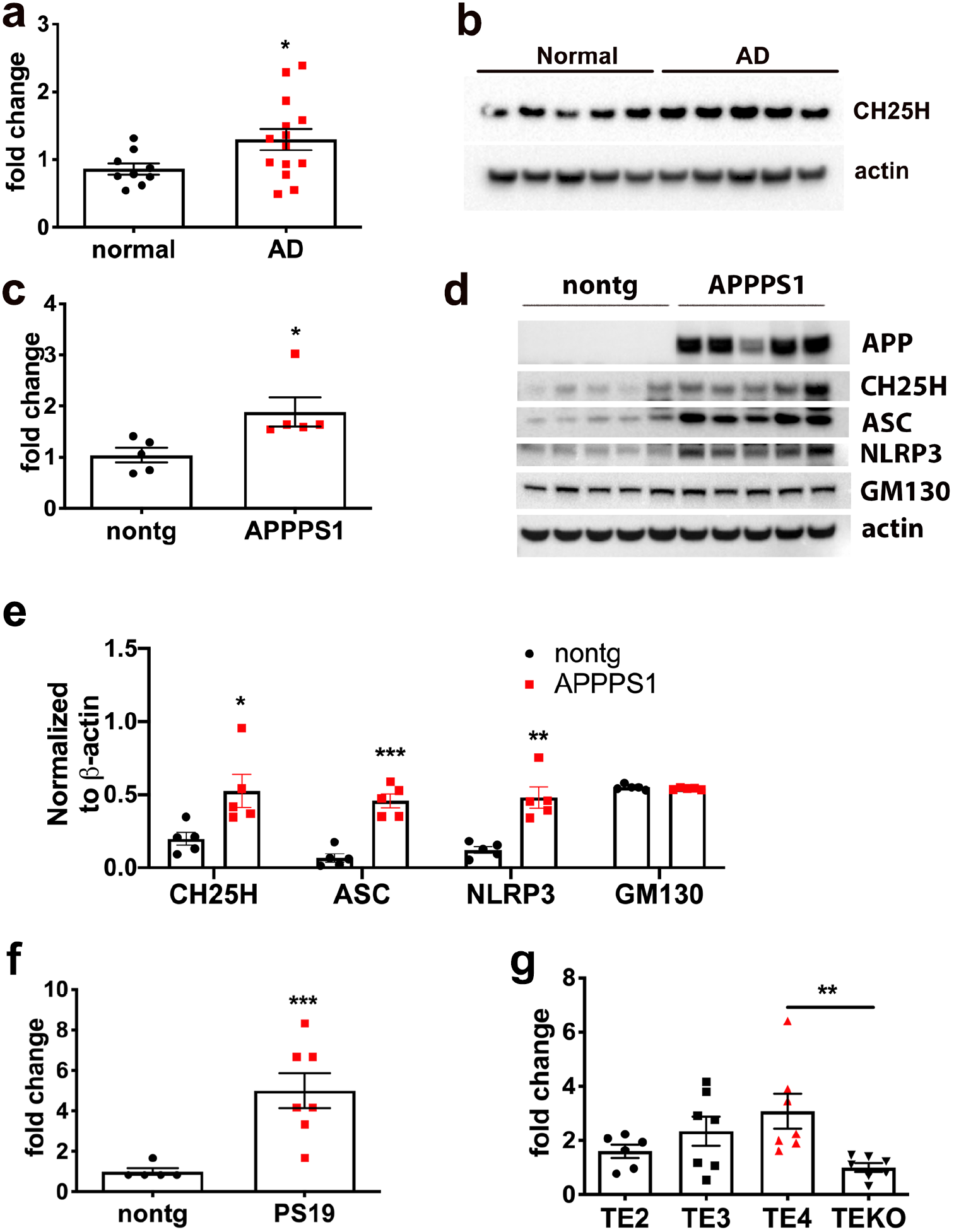
CH25H expression is increased in AD brain and AD transgenic mouse brain bearing amyloid or tau pathology. (a, b) Expression of CH25H at mRNA (a) or protein (b) levels in brain tissue of AD patients vs age-matched non-demented controls. (c,d) Expression of CH25H at mRNA (c) or protein (d) in APPPS1 transgenic mouse brain vs. age-matched non-tg littermates. (e) Quantification of d showing protein levels for CH25H, ASC and NLRP3 by normalization to β-actin. f) Expression of CH25H mRNA in PS19 tau P301S transgenic mouse brain vs non-tg littermates. g) Expression of CH25H mRNA in TE2, TE3, TE4 and TEKO mouse brain. Statistical significance was determined by student *t-test* with *\*p<0.05, ** p<0.01* or \*\*\**p<0.005* in a, c, e and f, or by Ordinary one-way *ANOVA* with Dunnett’s multiple comparisons test \**\*p<0.01* in g.

### LPS stimulates 25-HC production and CH25H expression in primary microglia

In macrophages, the TLR4 agonist lipopolysaccharide (LPS) stimulates expression of CH25H and production of 25-HC (43). In the central nervous system, CH25H is mainly expressed in microglia, the counterpart of peripheral macrophages, with very limited expression, if any, in other brain cell types, based on the Stanford transcriptome database generated by Barres group (http://www.brainrnaseq.org) (Supplemental Fig.1a). To explore a potential role for CH25H and its oxysterol product 25-HC in microglia-mediated innate immunity, we first measured 25-HC production by LC/MS in cultured microglia isolated from brain tissue of neonatal wild type mice in response to stimulation by LPS. A time- and dose-dependent increase of 25-HC production was observed in the cell lysate and medium of LPS-treated microglia compared to untreated microglia (Fig. 2a, b). As measured by qPCR, LPS stimulated the expression of the pro-inflammatory cytokines IL-1 β and TNFα as well as inflammasome genes such as NLRP3. It also potently upregulated CH25H mRNA in microglia (≥50 fold) (Fig. 2c). The increase in CH25H expression induced by LPS was further confirmed by Western blot using a CH25H specific antibody (Fig. 2c, insert). We next evaluated the effects of LPS on CH25H expression in the mouse *in vivo*. Wild type mice were treated with LPS (8.2mg/kg via i.p.) for 24 hrs, a marked increase in CH25H mRNA was detected in the hippocampus and cerebral cortex of LPS-treated mice compared to vehicle-treated mice (Fig. 2d). In contrast, the expression of CYP27a1 or CYP7b1 (two other enzymes involved in the cholesterol:oxysterol metabolic pathway) were not influenced by LPS treatment (Fig. 2d), suggesting that the induction of CH25H by LPS was highly specific. These results demonstrate that the production of 25-HC and the expression of CH25H are highly responsive to TLR4 stimulation in cultured primary microglia as well as in mouse brain *in vivo*.

**Figure 2:**
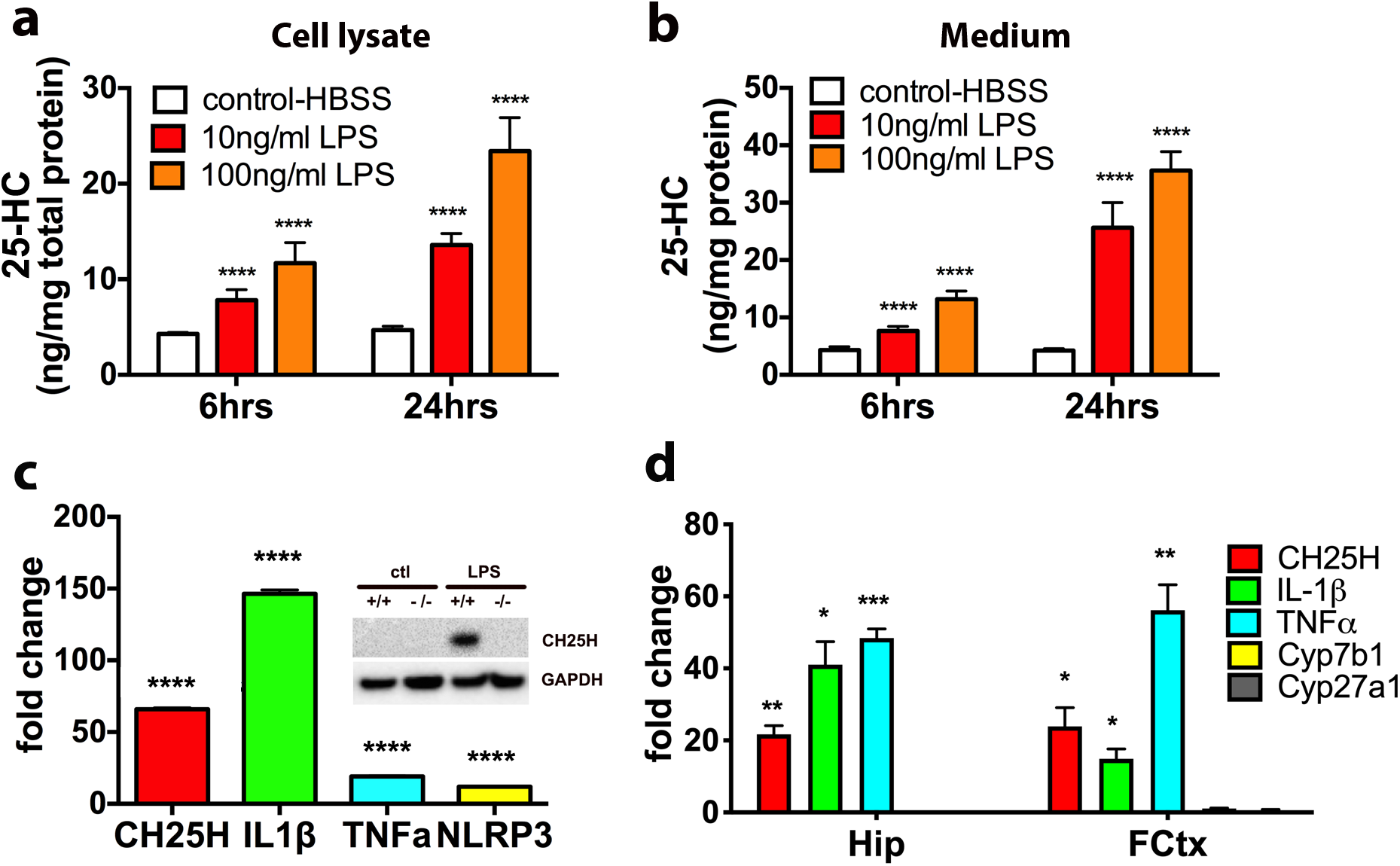
LPS stimulates 25HC production and CH25H expression in primary microglia and in mouse brain. a) LPS stimulates 25-HC production and secretion in primary microglia in a time-and dose-dependent manner. Primary microglia were treated with LPS (0, 10 and 100ng/ml) for 6 and 24 hours. The levels of 25-HC in cells (a) and media (b) were determined by GC-MS. *\*\*\*\*p<0.001* by ordinary one-way *ANOVA*. c) LPS induces the expression of CH25H, IL-1β, TNFα and NLRP3 inflammasome mRNA in primary microglia. The comparative gene expressions were determined by qPCR using RNA extracted from primary microglia with or without 10ng/ml LPS treatment for 24hrs. Insert: CH25H protein level in wt or CH25H-/-primary microglia treated with or without 10ng/ml LPS. It is a representative result of two independent experiments. d) Gene expression analysis of CH25H, IL-1β, TNFα, Cyp7b and Cyp27a1 in brain tissue of C57BL6 mice treated with 8.2mg/kg LPS (n=3) for 24 hours as determined by qPCR. * *p<0.05,**p<0.01 ***p<0.005* by student *t-test* comparing LPS-treated mouse brain to saline treated control brain.

### Depletion of 25-HC selectively attenuates LPS-induced IL-1β expression in primary microglia

To examine whether 25-HC is involved in the inflammatory response of microglia, we eliminated 25-HC production using microglia prepared from CH25H knockout (KO) mice (supplementary Fig. 1b). When WT or CH25H KO microglia were treated with LPS, we observed a significant reduction in the level of IL-1β secreted into the medium of CH25H KO microglia compared to WT microglia (Fig. 3a). The levels of IL-1α, a cytokine often co-released with IL-1β, were also reduced (Fig. 3b). In contrast, the production of TNFα (Fig. 3c) or IL-6 (not shown) were similar in both WT and CH25H KO cells treated with LPS. The addition of 25-HC to CH25H KO microglia fully rescued the attenuated IL-1β/α production observed in CH25H KO microglia to a comparable level as WT microglia (Fig. 3d). These data suggest that 25-HC contributes to the LPS-triggered IL-1β production by microglia. To directly evaluate the effect of 25-HC on IL-1β/α production, we treated WT microglia with 25-HC alone or in combination with LPS. Compared to LPS treatment alone, the addition of 25-HC in the presence of LPS resulted in a marked dose-dependent increase of microglial IL-1β and IL-1α secretion while 25-HC treatment alone had no effect (Fig. 3e).

**Figure 3:**
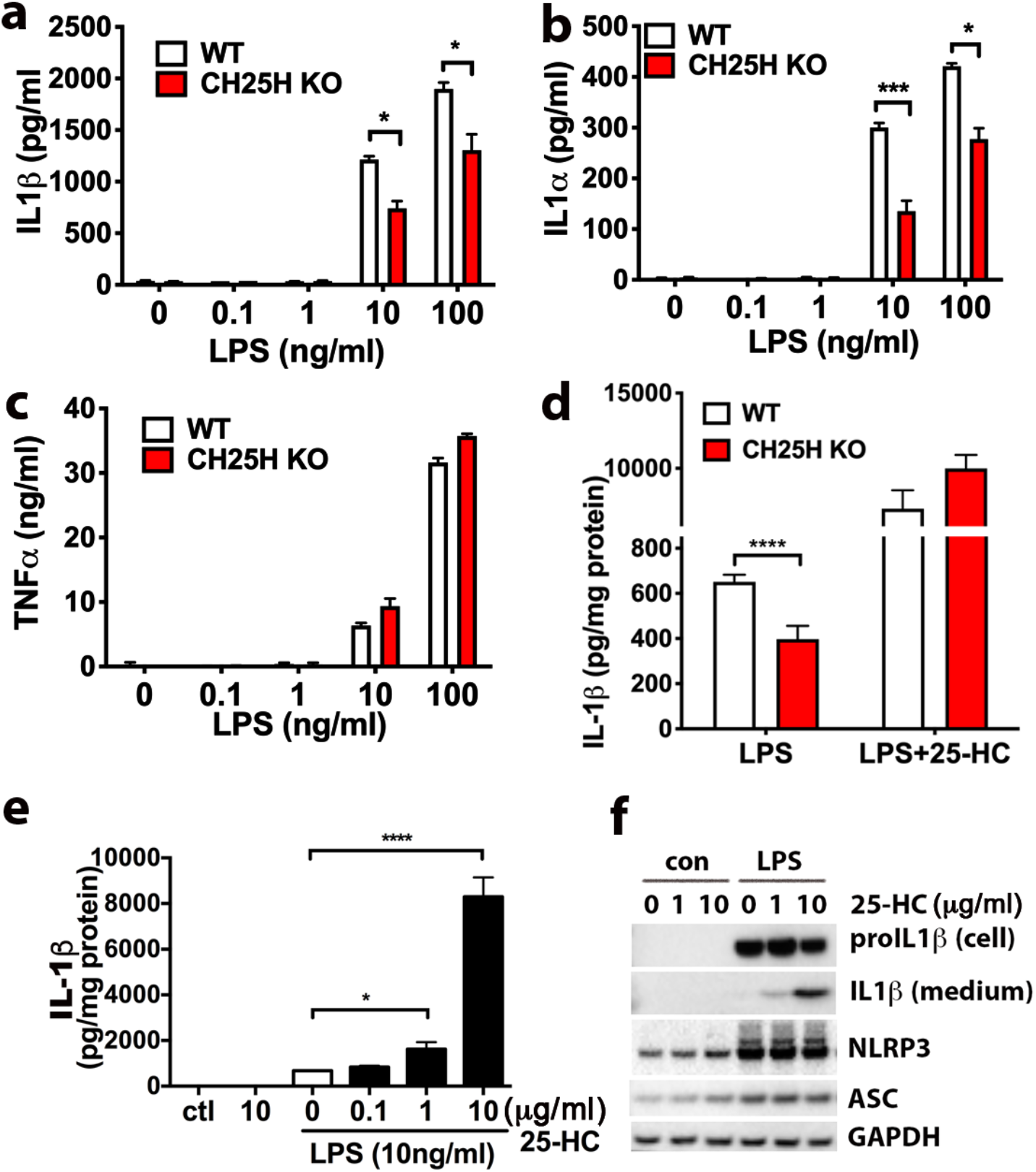
25-HC selectively amplifies LPS-induced IL-1 β expression and secretion. a-c) WT and CH25H KO primary microglia were treated with LPS (0, 0.1, 1, 10, 100ng/ml) for 24hrs. The levels of secreted IL-1β(a), IL-1α (b) and TNFα (c) in the medium were measured by ELISA. d) The levels of secreted IL1βfrom WT and CH25H KO microglia treated with LPS (10ng/ml) with or without 25-HC (10μg/ml) were measured by ELISA. e) Primary microglia were treated with 10ng/ml LPS in the presence of different concentrations of 25-HC for 24 hours. The levels of IL-1βin the media were determined by ELISA and f) the levels of intracellular pro-IL1βand mature IL1βsecreted in the media as measured by Western blotting. Statistical analyses were determined by multiple *ttest* in a, b, c, d; one-way *ANOVA* in e. * *p<0.05, **p<0.01, ***p<0.005, ****p<0.001*. The data shown are representative for three or more independent experiments.

Mature IL-1β (17kDa) is produced from its 31 kDa pro-IL-1β by the inflammasome complex and rapidly secreted into medium. We next examined the effects of 25-HC on the level of pro-IL-1β protein remaining in cells and mature IL-1β protein released into the medium by Western blotting. LPS treatment markedly increased the cellular level of pro-IL-1β as well as the inflammasome proteins NLRP3 and ASC1, resulting in a limited amount of 17kDa IL-1β produced and secreted into the medium. However, the addition of 25-HC markedly and dose-dependently stimulated the release of active 17kDa IL-1βinto the medium (Fig. 3f). The intracellular protein levels of unprocessed pro-IL-1β, NLRP3 or ASC were not influenced by the presence of 25-HC (Fig. 3f). Therefore, 25-HC may regulate IL-1β production at a posttranslational level. Together, these results suggest that 25-HC modulates LPS-activated inflammatory responses by selectively promoting mature IL-1β production.

### APOE4-expressing microglia show exaggerated IL-1β production in response to LPS and 25-HC treatment

Previous studies have shown that APOE isoforms differentially influence the innate immune response of microglia (26, 27). We therefore examined the effects of the common APOE isoforms on both LPS and 25-HC-enhanced production of IL-1β in microglia. Microglia were prepared from neonatal mice expressing human APOE2 (E2), APOE3 (E3), or APOE4 (E4) at the mouse APOE locus (56–58). Consistent with previous reports, E4-expressing microglia produced higher levels of IL-1β than E2-expressing cells or APOE deficient cells (EKO) after 6 h (Fig. 4a) or 24 h (Fig. 4b) following LPS treatment alone. As expected, 25-HC dose-dependently increased IL-1β production at 6 h (Fig. 4a) and at 24 h (Fig. 4b). Strikingly, co-incubation with 25-HC resulted in a marked potentiation of IL-1β production in E4-expressing microglia compared to E2-expressing or EKO microglia at each concentration of 25-HC tested (Fig. 4a and 4b), resulting in significantly higher levels of IL-1β production from E4 microglia than that from E2 or EKO microglia (Fig. 4a and 4b). Although LPS induced greater IL-6 production in E4-expressing microglia, 25-HC treatment did not influence the production of IL-6 (Fig. 4c). We further compared the IL-1β-inducing activity of 25-HC between E4 and E3 microglia. A higher amount of secreted (extracellular) IL-1β was observed in E4 microglia than in E3 microglia treated with both LPS and 25-HC (Fig. 4d). Consistently we detected more mature IL-1β protein (17kd) in the medium of E4 microglia than in the medium of E3 microglia, while the levels of intracellular pro-IL-1β did not increase in cells treated with 25-HC (Fig. 4e). Together, these data demonstrate that apoE isoforms differentially influence the ability of 25-HC to augment the secretion of IL-1β production in LPS-activated microglia and the presence of APOE4 markedly augments the effects of 25-HC in promoting IL-1β production, shifting the dose-response for 25-HC substantially to the left. Lastly, the production of 25-HC by E2 or E4-expressing microglia was measured. We also found that E4 microglia produced greater amount of 25-HC measured in both cells and medium than E2 microglia when treated with LPS (Fig. 4f).

**Figure 4:**
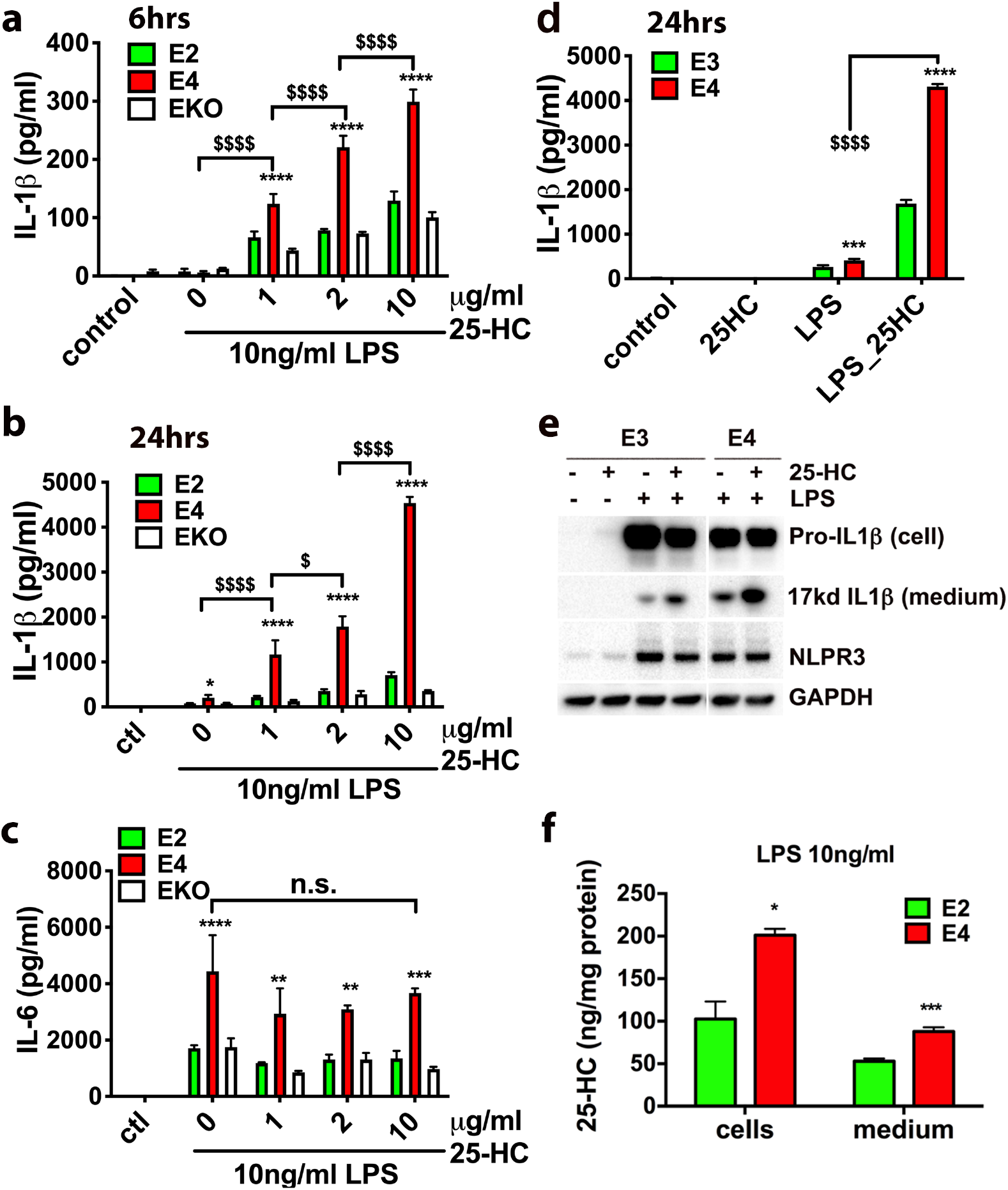
Exaggerated IL-1β and 25-HC production in LPS-activated microglia expressing human ApoE4. The levels of IL-1βor IL-6 secreted into the medium in apoE2-or apoE4-expressing microglia or apoE KO microglia treated with LPS (10ng/ml) and 25-HC (0, 1, 2, or 10μg/ml) for 6 h (a) or 24 h (b and c). The levels of IL-1βsecreted in the medium of apoE3-or apoE4-expressing microglia after 24 h treatment with LPS (10ng/ml) and 25-HC (10μg/ml) (d, e) and the levels of 25-HC in these cells or medium were determined by GC-MS (f). Statistical significances were determined by two-way *ANOVA* with multiple comparisons in a, b, c, d or unpaired student *t-test* in f. * *p<0.05, **p<0.01, ***p<0.005, ****p<0.001, respectively.* The data shown are representative for three or more independent experiments.

### Augmentation of LPS-induced IL-1β induction by 25-HC is enantioselective

To examine the specificity of 25-HC, we first tested the effects of both the 25-HC precursor cholesterol and another cholesterol metabolite 7α-HC on IL-1β production. Comparing to the promoting effects of 25-HC on IL-1β/α production, coincubation of cholesterol or 7α-HC with LPS at a similar concentration as 25-HC did not promote LPS-induced IL-1β/α production in microglia (Fig. 5a, b, c). We further evaluated the IL-1βinducing activity of *ent*-25-HC, the inactive enantiomer of 25-HC (59), and found that ent-25HC exhibited only very weak IL-1β-inducing activity and was at least an order of magnitude less potent than 25-HC (Fig. 5d). These results demonstrate that the IL-1β induction by 25-HC is enantioselective and thus likely mediated via a specific protein target(s).

**Figure 5:**
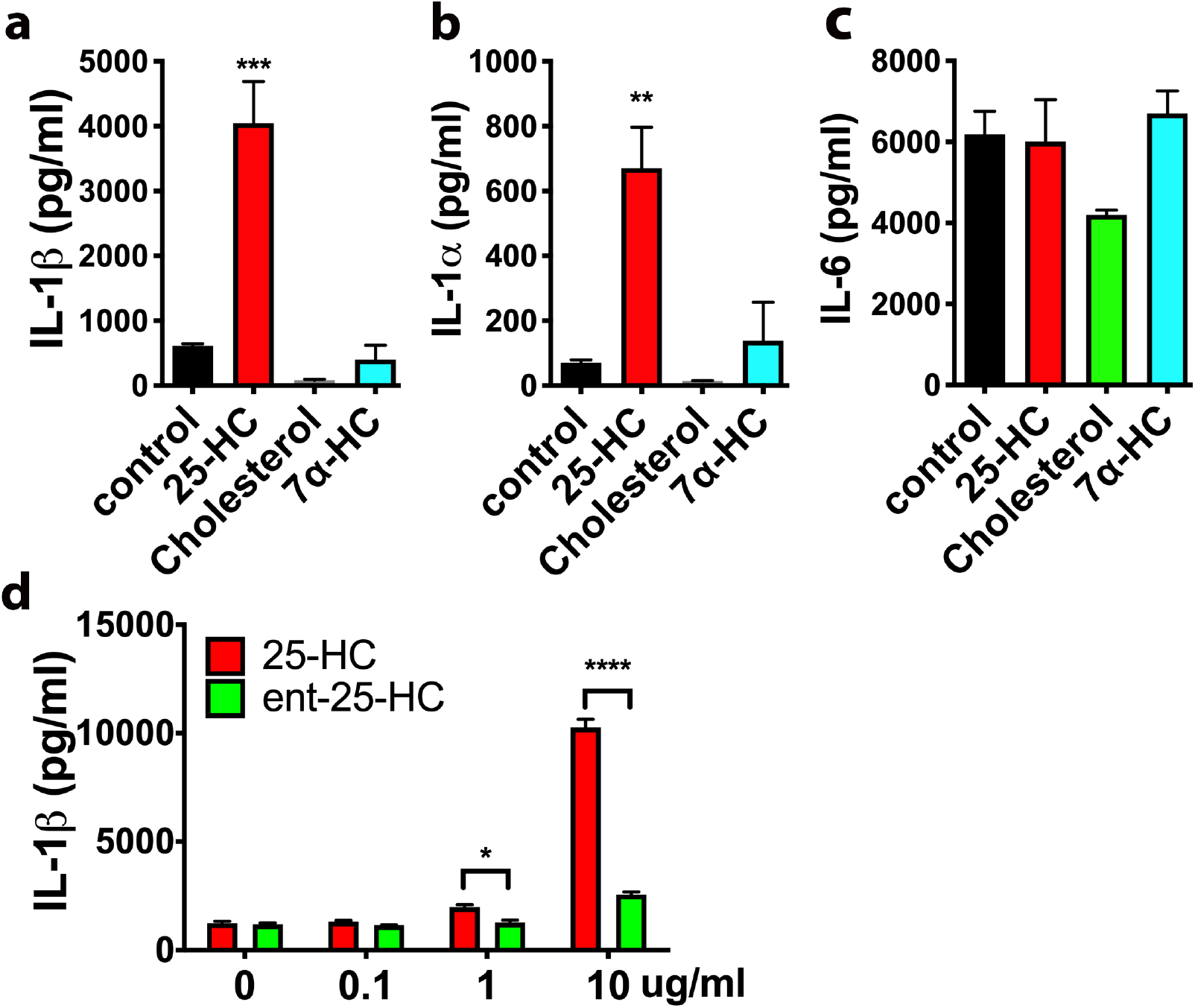
IL-1β/α induction by 25-HC is highly specific. The levels of secreted IL-1β(a), IL-1α (b) or IL-6 (c) in the medium of primary microglia treated with LPS (10ng/ml) in the presence of 25-HC (10μg/ml), Cholesterol (10μg/ml) or 7α-HC (10μg/ml) for 24 hrs. d) Ent-25-HC (10μg/ml) has much weaker effects in augmenting IL-1βproduction in primary microglia treated with LPS (10ng/ml) for 24hr. The levels of cytokines were determined by ELISA. Statistical significances were determined by ordinary two-way *ANOVA* with Tukey multiple comparisons test. * *p<0.05, **p<0.01, ***p<0.005, ****p<0.001.* The data shown are representative for two or more independent experiments.

### 25-HC induces IL-1β via activation of caspase-1 and the inflammasome

Active 17kD IL-1β is produced from pro-IL-1β after proteolytic cleavage by caspase-1. Formation of ASC, recognized as large perinuclear cellular aggregates, is a hallmark of inflammasome activation that correlates with caspase-1 cleavage and release of mature IL-1β (60). To further address if 25-HC activates the inflammasome in microglia, we compared the number of cells with adaptor protein apoptosis associated speck-like protein containing a CARD (ASC) speck in microglia treated with LPS alone or LPS combined with 25-HC. The number of ASC speck-containing cells significantly increased following treatment with LPS and 25-HC compared to LPS alone (Fig. 6a). 25-HC treatment alone, however, did not induce ASC speck formation (Fig. 6a, b). We further found that the induction of ASC speck by LPS and 25-HC is dependent on TLR4, because no ASC speck formation was detected in TLR4 KO microglia treated with LPS and 25-HC (Fig. 6c, d). The induction of IL-1β by LPS and 25-HC was also markedly reduced or eliminated in TLR4 KO microglia (Fig. 6c, d). These data suggest that 25-HC augments IL-1β secretion via activation of the inflammasome in a TLR4-dependent manner. Activation of NLRP3 inflammasome triggers oligomerization of caspase-1 that cleaves pro-IL-1β to biologically active IL-1β. To examine if the induction of IL-1β by 25-HC is caspase-1-dependent, primary microglia were treated with LPS and 25-HC in the presence of VX765, a prodrug of VRT-043198 that selectively inhibits the caspase-1 subfamily of cysteine proteases (61). Treatment with VX765 completely inhibited the effect of 25-HC on IL-1β production (Fig. 6e), suggesting that 25-HC induces IL-1βproduction by activating the inflammasome and caspase-1. Potassium efflux is one of the common mediators of NLRP3 inflammasome activation in response to diverse stimuli (15). When potassium efflux was blocked by a high concentration of extracellular KCl, we found that the induction of IL-1β by LPS and 25-HC was effectively prevented by 50mM KCl (Fig. 6f). This result confirms that activation of the inflammasome by LPS is augmented by 25-HC and further suggests that 25-HC regulates IL-1β induction upstream of potassium efflux.

**Figure 6:**
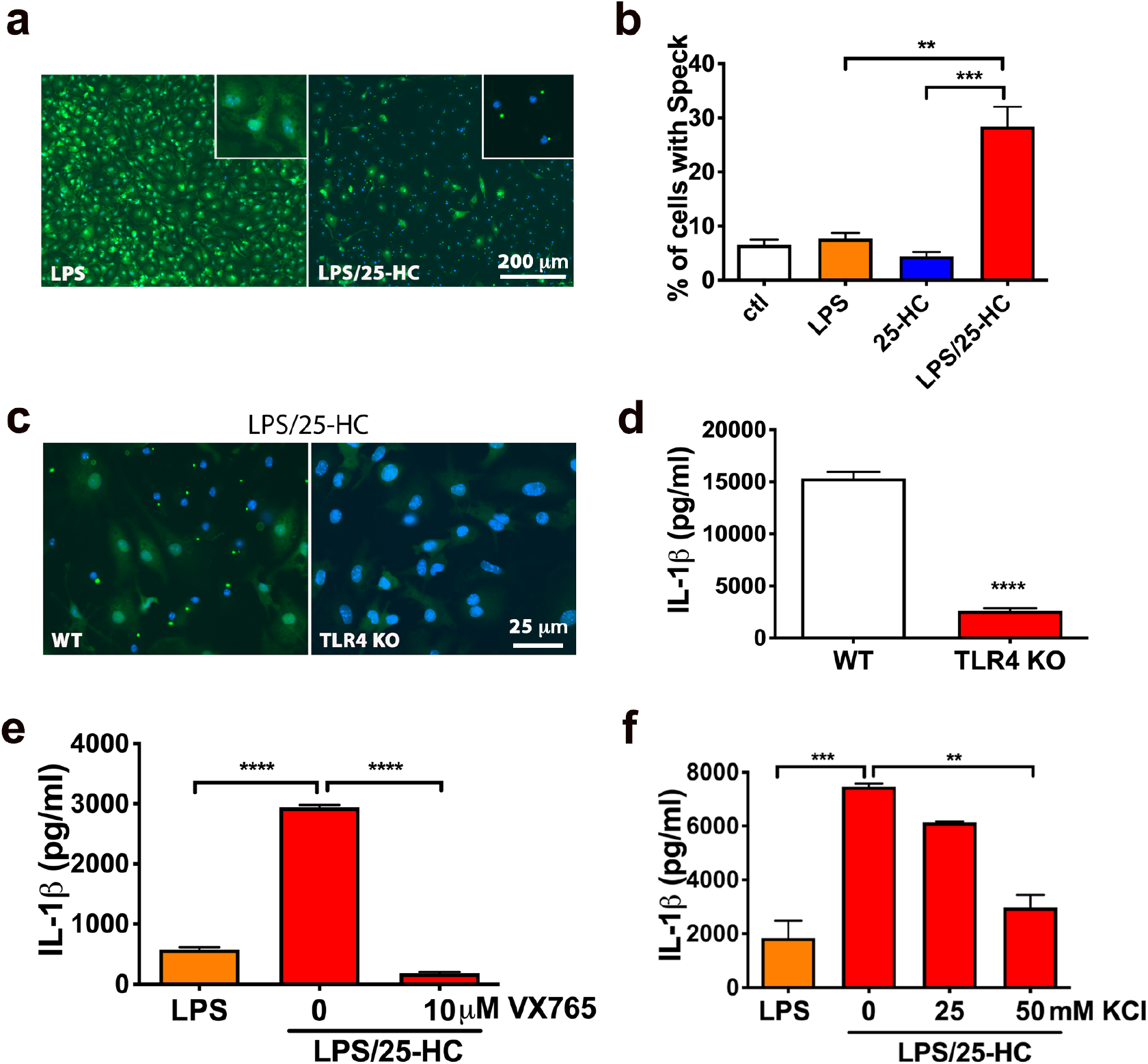
25-HC induces inflammasome activation. ASC specks in microglia treated with LPS (10ng/ml) w/o or with 25-HC (10μg/ml) were stained by ASC antibody (green) and DAPI for Nuclei (blue) (a). Quantification of ASC specks in microglia treated with medium alone (ctl), medium containing LPS (10ng/ml), 25-HC (10μg/ml) or LPS (10ng/ml) plus 25-HC (10μg/ml) (b). WT or TLR4 deficient microglia were treated with LPS (10ng/ml) and 25-HC (10μg/ml) for 24 h followed by ASC antibody staining (green) and DAPI (blue) (c) or ELISA measurements of secreted IL1β(d). Inhibition of caspase 1 by VX765 (e) or a high concentration of potassium (50mM) (f) in the medium prevents 25-HC-dependent IL-1βproduction in microglia treated with 10ng/ml LPS for 24 h. Statistical analyses were determined by ordinary two-way *ANOVA* with Tukey multiple comparisons test in c, e and f or student *ttest* in d. * *p<0.05, **p<0.01, ***p<0.005, ****p<0.001.* The data shown are representative for three or more independent experiments.

## Discussion

### CH25H and 25-HC in innate immunity

25-hydroxycholesterol (25-HC) is an enzymatically derived oxidation product of cholesterol which is produced primarily by circulating and tissue-resident macrophages and which has been reported to have both anti-inflammatory as well as proinflammatory effects in various model systems of innate immunity (44). The enzyme cholesterol-25-hydroxylase (CH25H) which catalyzes the synthesis of 25-HC from cholesterol is markedly upregulated in macrophages following stimulation with interferon and the TLR4 ligand, LPS (43). 25-HC has also been reported to regulate cholesterol metabolism by suppressing cholesterol biosynthesis via SREBP processing and facilitating reverse cholesterol transport via activation of liver X receptors (LXRs) and various downstream genes (62). 25-HC has been shown to be a potent antiviral oxysterol and likely mediates the antiviral action of interferons against a variety of enveloped DNA and RNA viruses (44, 63). Although 25-HC’s anti-inflammatory actions have been widely documented (see below), it has also been reported to have proinflammatory effects and may contribute to the tissue damage in mice observed following influenza infection by acting as an amplifier of inflammation by macrophages via an AP-1-mediated mechanism (64). CH25H deficient mice have also been reported to show decreased inflammatory-mediated pathology and death following influenza infection (64), reduced immune responses following experimental autoimmune encephalomyelitis (EAE)(65) and in a mouse model of X-linked adrenoleukodystrophy (X-ALD) (66), again supporting a proinflammatory and potentially “toxic” function of 25-HC. Moreover, 25-HC was recently identified as an integrin ligand and to directly induce a proinflammatory response in macrophages (67).

In our study we show that CH25H is expressed in microglia *in vitro* and further demonstrate that the TLR4 agonist LPS induces a marked upregulation of CH25H expression and 25-HC production and secretion. This increase in CH25H expression and 25-HC production in microglia was accompanied by corresponding increases in the secretion of the inflammatory cytokines IL-1β, IL-1α, and TNFα. Reductions in both LPS-stimulated IL-1β and IL-1α secretion (but not TNFα secretion) were observed in CH25H-deficient microglia, suggesting an autocrine or paracrine effect of 25-HC in amplifying proinflammatory signaling in microglia (see below). Treatment of CH25H-deficient microglia with 25-HC restored the effect of LPS on IL-1β/α secretion. We also observed an increase in CH25H mRNA following LPS treatment of wild-type mice *in vivo*, consistent with the *in vitro* microglia data.

### Possible roles of CH25H and 25-HC in AD

CH25H is located on chromosome 10q23, a region strongly linked to AD (47). In a large scale AlzGene meta-analysis including 1282 AD patients and 1312 controls from five independent populations (French, Russian, USA, Swiss, Mediterranean), a significant association of rs13500 &#8216;T’ allele and haplotypes in CH25H promoter were found to be significantly associated with the risk of developing AD and with different rates of Aβ/amyloid deposition (47). Although the association of this rs13500 promoter polymorphism was not found in two sequent studies (68, 69), a recently study carried out in a Turkish cohort of AD subjects and controls also revealed a strong association of CH25H rs13500 and AD, and an even stronger risk factor in the presence of *APOE4* (70). More recently, several genome-wide expression studies carried out in models of accelerated aging, AD pathology and neuroinflammation have all identified CH25H as being significantly upregulated in brain (51–53). Here we also show that CH25H is upregulated in AD brain tissue compared to age-matched controls as well as in two mouse models of AD pathology; APPPS1 transgenic mice, tau transgenic mice and a recently described APOE4xP301S (TE4) tau transgenic mouse model of accelerated tau pathology and neurodegeneration (37). These findings suggest that 25-HC may be involved in AD pathogenesis, especially given its reported proinflammatory properties and our data on its marked potentiation of cytokine expression and secretion from microglia stimulated by the TRL4 agonist, LPS.

### CH25H, 25-HC and APOE genotype

Given the important role of APOE4 as a genetic risk factor for AD and its reported role in regulating innate immunity in brain (71), we examined whether CH25H expression and 25-HC production in microglia were impacted by APOE genotype. First, we found that apoE4-expressing microglia produced significantly more 25-HC in response to LPS treatment than apoE2-expressing microglia. We also found that apoE4-expressing microglia produced more IL-1β and IL-6 in response to LPS treatment as has been previously reported (27). To our surprise, co-incubation of 25-HC with LPS markedly augmented IL-1β production in apoE4-expressing microglia compared to either apoE2-expressing microglia or apoE deficient (knockout) microglia, markedly shifting the dose-response curve for 25-HC to the left. In fact, relatively low concentrations of 25-HC (~1uM) stimulated IL-1βproduction in apoE4-expressing (vs. apoE2-expressing) microglia, again demonstrating that 25-HC’s proinflammatory effects in this in vitro model of innate immunity are APOE isoform-dependent. Previous work has shown that treatment with LPS induces higher levels of various cytokines (including IL-1β) in the serum of human APOE4 carriers than APOE3 homozygotes (26) and in the brains of apoE4-expressing targeted replacement mice (27). *In vitro*, apoE4-expressing microglia exhibit higher “innate immune reactivity” following LPS treatment measured by both cytokine and NO production (27). Moreover, APOE genotype alters glial activation in response to LPS treatment (72). Together, with our *in vivo* data in several AD mouse models demonstrating higher brain levels of microglial and brain CH25H mRNA, we hypothesize that 25-HC may be an important proinflammatory chemical messenger whose production and secretion will greatly amplify cytokine secretion in apoE4-expressing microglia in a paracrine or autocrine manner, and may thus contribute either indirectly or even directly to the neuroinflammation and neurodegeneration that characterize AD. In this regard, Jang and colleagues (66) have recently shown that 25-HC has proinflammatory actions in a study of X-linked adrenoleukodystrophy (X-ALD), a progressive neurodegenerative disorder characterized by accumulation of very long-chain fatty acids (VLCFA). They observed that 25-HC is markedly increased in X-ALD brain tissue, promotes IL-1β production and neuroinflammation and is directly neurotoxic when administrated to brain *in vivo* (66).

### 25-HC, IL-1β production and inflammasome activation

Consistent with the work of Jang et al (66), we provide evidence supporting a proinflammatory role of 25-HC in microglia by promoting mature 17kD IL-1β production via inflammasome activation. However, we did not observe any change of pro-IL-1βmRNA or protein levels in 25-HC treated microglia, suggesting that 25-HC augments cytokine production via a posttranslational mechanism. The induction of IL-1βproduction is dependent on two signals: first, activation of TLR4 on the cell surface by stimuli such as LPS leading to IL-1βmRNA generation and pro-IL-1βproduction. A second process derives from inflammasome activation by stimuli such as ATP which leads to recruitment and activation of caspase-1, a protease that cleaves pro-IL-1β into mature IL-1β. We found that 25-HC efficiently promotes IL-1β production in the presence of LPS, however, 25-HC does not activate IL-1β production by itself at either the mRNA or protein level. These observations suggest that 25-HC might directly or perhaps indirectly activate caspase-1 inflammasome activity in microglia. In fact, we observed markedly reduced IL-1β production when 25-HC and LPS were coincubated in the presence of the caspase-1 inhibitor VX765 or when K+ efflux was blocked by high concentrations of extracellular K+. Together, these data suggest that augmentation of inflammasome activity and IL-1β production by 25-HC occurs post-translationally upstream of K+ efflux. Our observations, together with Jang et al. (66), are not consistent with the previous report by Reboldi et al. (73). In activated BMDMs, they found that low concentration of 25-HC inhibited IL-1β production and CH25H deficiency caused augmented transcription and secretion of the cytokine IL-1β. They also showed that 25-HC regulates IL-1β production via repressing SREBP-mediated transcription (73). Following this, Dang et al. later showed that up-regulating CH25H and 25-HC production reduce inflammasome activity and IL-1β levels in LPS-activated macrophages (74). The discrepancy between our data may result from treatment conditions (such as LPS or 25HC concentrations, treatment duration time etc.) and the different cell types used in our respective experiments.

With advances in genomic sequencing and bioinformatics, more genetic risk factors and related molecular pathways have been identified as potentially important in the etiology and pathogenesis of AD. These risk genes associated with late onset-AD (LOAD) point to both lipid metabolism and immune mechanisms as contributing to AD pathology. However, how the components of two distinct essential cellular pathways are connected to clinical and pathological disease phenotypes and finally contribute to the neurodegeneration in AD patients remains unclear. Our present study has identified an interaction among APOE genotype, cholesterol metabolism to the oxysterol 25-HC and the cytokine IL-1β in microglia. Our data suggest that microglial expression and activation of CH25H and consequent 25-HC production may be an important mediator of the progressive neuroinflammation that characterizes neurodegenerative disorders like AD. Importantly, the proinflammatory effects of 25-HC we observe in primary microglia are APOE isoform-dependent as apoE4-expressing microglia secrete more 25-HC and are markedly more sensitive to the proinflammatory actions of 25-HC than apoE2 or apoE3-expressing microglia. Thus, the immune oxysterol 25-HC may play an important role in the pathogenesis, i.e. the neuroinflammation and neurodegeneration, that characterize AD and perhaps other neurodegenerative disorders.

## Materials and Methods

### Ethics Statement

All experiments were conducted in accordance with relevant NIH guidelines and regulations related to the Care and Use of Laboratory Animals and human tissue. Animal procedures were performed according to protocols approved by the Research Animal Resource Center at Weill Cornell Medicine.

### Animals

APPPS1-21 transgenic mouse model (54) co-expressing human APP KM670/671NL and Presenilin 1 L166P under the control of a neuron-specific Thy1 promoter element was kindly provided by Dr. Mathias Jucker through an agreement with Koesler. These mice were intercrossed and maintained on a C57BL/6J background. PS19 expressing human P301S tau under PrP promotor were purchased from Jackson laboratory (#008169) and backcrossed and maintained on a C57BL/6 background. CH25H knockout mice (75) were purchased from the Jackson laboratory (JAX stock #016263) and maintained as homozygotes. Human APOE targeted replacement mice with the human APOE2, APOE3 or APOE4 coding sequences inserted behind the endogenous murine APOE promoter on a C57BL/6 J background were provided by P.M. Sullivan of Duke University (56–58). APOE-/-mice were purchased from Taconic. P301S tau transgenic mice that are homozygous for human APOE2 (TE2), APOE3 (TE3), APOE4 (TE4) or with no expression of apoE (TEKO) (C57BL/6) were generated by the Holtzman laboratory at Washington University, St. Louis as described previously (37). TLR4 knockout mice were purchased from the Jackson laboratory (JAX stock #029051) and maintained as homozygotes. All animals were maintained in a pathogen-free environment, and experiments on mice were conducted according to the protocol approved by the Weill Cornell Medicine Animal Care Committee.

### Human brain specimens

Frontal cortical tissue samples from AD patients or age-matched controls with no reported clinical signs of dementia (≥80yrs) were obtained from the Brain Bank of the University of Miami Miller School of Medicine, the Human Brain and Spinal Fluid Resource Center of the Greater Los Angeles VA Healthcare System at the West Los Angeles Healthcare Center, University of Maryland Brain and Tissue Bank, and the New York Brain Bank at Columbia University through requests from the NIH NeuroBioBank. All procedures were approved by the Weill Cornell Medicine Human Biology Research Ethics Committee.

### Culture and treatment of primary microglia

Primary neonatal microglia were prepared from cerebral cortices of 1-3 day old neonatal mice as previously described (76). Cell suspensions of cerebral cortices were seeded into a 75 ml flask and cultured in DMEM/F12 medium containing 10% FBS and 5ng/ml GM-CSF. Microglial cells floating on top of the astrocyte layer were harvested at 12 DIV by shaking for 2 hrs at 200 rpm and seeded onto 48 well (3×10^5^/well) or 24 well (6×10^5^/well) culture plate in DMEM/ F12/10%FBS medium without GM-CSF. Over 98% of the cells were determined to be microglia (Iba-1 positive) by immunohistochemistry. After seeding for 24 hrs, cells were washed once with serum-free medium and treated with various reagents in serum-free DMEM/F12 medium supplemented with 0.02% BSA. The reagents used in microglia treatment were: LPS (Sigma, L5293, Escherichia coli, 0111:B4); ATP (sigma A2383); 25-hydroxycholesterol (Avanti#700019 or Sigma H1015); cholesterol (Avanti#700100); 7 α-hydroxycholesterol (Avanti#700034); VX-765 (Medchemexpress). Ent-25-hydroxycholesterol was synthesized as described (77).

### Cytokine ELISAs

Supernatants from cell cultures were collected and the concentrations of IL-1β(BioLegend#432601), IL-1α (Biolegend#433401), IL-6 (Bon Opus Biosciences#BE010059B), and TNFα (Biolegend#430901) were determined by ELISA according to the manufacturer’s instructions. All cytokine levels were normalized to microglial protein levels determined by BCA assay.

### ASC speck analysis

For measuring ASC speck formation, mouse primary microglia were seeded at 0.15×106/well in 8-well chamber Millicell EZ slides (Millipore PEZGS0816) and allowed to attach overnight. The following day, the cells were treated with 100ng/ml LPS in the presence of absence of 10μg/ml 25-HC over 16hrs. The cells were fixed in 4% paraformaldehyde and then washed three times in PBS with Tween 20 (PBST). After permeabilization with Triton X-100 and blocking with 10% bovine serum albumin in PBS, the cells were incubated with anti-mouse ASC antibody (Cell Signaling#67824) overnight at 4°C. After washing with PBST, the cells were incubated with secondary antibodies (Jackson ImmunoResearch) in PBS for 30 min and rinsed in PBST. The slides were mounted with mounting solution containing DAPI. Images were taken using a Nikon *eclipse* 80i microscope. For each treatment condition, 3-5 pictures taken from different areas in the well at 20x magnification were used for counting cells containing ASC speck. The total number of cells was determined by visualizing DAPI positive nuclei. Each experimental condition was repeated more than three times.

### Immunoblotting

To detect CH25H protein, microsomal membranes were prepared as described previously (75, 78), solubilized in a small volume of buffer A (50 mM Tris-Cl, pH 7.4, 1mM EDTA, 0.05% (w/v) SDS), mixed with an equal amount of HMG-CoA solubilization buffer (62.5 mM Tris-Cl, pH6.8, 15% SDS, 8 M urea, 10% glycerol, 100 mM dithiothreitol). 100 μg lysate was incubated with NuPAGE LDS sample buffer at 37C for 20min followed with separation by NuPAGE 4-12% Bis-Tris gel and transferring to nitrocellulose membrane (Amersham Biosciences). For other proteins, cell lysates (~40μg of protein/lane) were resolved in 4-20% Bis-Tris gels and transferred to nitrocellulose membranes. Blots were incubated with antibodies at 4°C overnight followed by horseradish peroxidase-coupled secondary antibodies and ECL developing kits (Amersham Biosciences). The images were taken using Bio-Rad Molecular-Imager ChemiDoc XRS+ and densitometry of the bands was measured with Bio-Rad Image lab software and all values were normalized to β-actin or glyceraldehyde-3-phosphate dehydrogenase (GAPDH). Antibodies used for immunoblotting were: mouse anti-human CH25H (hybridoma supernatant, neat, kindly provided by Dr. David Russell, University of Texas, Southwestern medical center)(75), mouse anti-GAPDH antibody (GeneTex, GT239), mouse-anti-β-actin (GeneTex, GT5512), mouse anti-human 6E10 for full length APP (Covance, SIG393206), rabbit anti-mouse ASC antibody (Cell Signaling#67824), mouse anti-NLRP3 (AdipoGen, Cryo2, AG-20B-0014-C100), mouse anti-GM130 (Santa Cruz, sc-55591), rabbit anti-IL-1β(Abcam, ab9722).

### Quantification of 25-hydroxycholesterol

Primary microglia were prepared and treated as described above. Media was collected and frozen at −80°C after removing floating cells. For each sample, 5 µL of methanol or 5 µL of deuterated internal standard at a concentration of 500 ng/mL were added to 50 µL of microglia growth media separately before being mixed and then hydrolyzed using 1N KOH at 90°C for two hours. The samples were then liquid-liquid extracted with methyl tert-butyl ether and the organic phase evaporated to dryness under air at 50°C. Sample residues were reconstituted in 100 µL of 80% methanol. Reconstituted samples (5ul) were then injected onto an Eksigent microLC 200 system. Separation was effected with a Waters Acquity 1 mm x 50 mm C18 reverse-phase column at 50 µL/min over seven minutes. Data were acquired by an ABSciex QTRAP 5500 mass spectrometer using the Turbo Spray source maintained at 300°C. Spray voltage was maintained at 4000 volts, curtain gas at 40 L/min, gas 1 at 30 L/min, and gas 2 at 30 L/min. Chromatographic peak areas of transition 385.4/367.4 (CE=25V, DP=60V) were integrated and quantified using MultiQuant 3.0 software (ABSciex).

### RNA Isolation, Real-time RT-PCR and nanostring analysis

Total RNA was isolated from primary microglia or mouse brain tissue with the PureLink RNA mini kit (Invitrogen#12183018A) and reverse transcribed to cDNA using SuperScript IV VILO Master Mix with ezDNase Enzyme (Thermo Fisher, # 11766050) following the manufacturer’s protocol. Quantitative real time PCR were performed using Taqman gene expression assays and gene expression master mix (Applied Biosystems, #4369016). The changes in gene expression were normalized to β-actin or glyceraldehyde-3-phosphate dehydrogenase (GAPDH).

### Statistical Analysis

Data are expressed as mean ± SEM. Significance was assessed with Student’s *t-*test, one-way or two-way *ANOVA* followed by Tukey multiple comparisons test or Bonferroni’s post hoc test using Prism version 8.0 software (GraphPad).

## Acknowledgements

We thank our colleagues in the Appel Alzheimer’s Disease Research Institute for assistance and suggestions. We thank Dr. Li Gan for her critical comments for this work.

## Authors’ contributions

SMP and WL conceived the project and designed the experiments. MYW, ML, JJD, YS, SMP and WL analyzed the data. MYW, SMP and WL wrote the paper. MYW, ML, YS, JD and WL performed all experiments, with help or guidance from PMS, DFC, DMH and GAP. MQ synthesized *ent-*25-HC as previously described(77). All authors read and commented on the manuscript.

## References

1. Heneka MT, Carson MJ, El Khoury J, Landreth GE, Brosseron F, Feinstein DL, et al. Neuroinflammation in Alzheimer’s disease. Lancet Neurol. 2015;14(4):388–405.

2. Heneka MT, Golenbock DT, Latz E. Innate immunity in Alzheimer’s disease. Nat Immunol. 2015;16(3):229–36.

3. Landreth GE, Reed-Geaghan EG. Toll-like receptors in Alzheimer’s disease. Curr Top Microbiol Immunol. 2009;336:137–53.

4. Gold M, El Khoury J. beta-amyloid, microglia, and the inflammasome in Alzheimer’s disease. Semin Immunopathol. 2015;37(6):607–11.

5. Walker DG, Lue LF, Beach TG. Gene expression profiling of amyloid beta peptide-stimulated human post-mortem brain microglia. Neurobiol Aging. 2001;22(6):957–66.

6. Akama KT, Van Eldik LJ. Beta-amyloid stimulation of inducible nitric-oxide synthase in astrocytes is interleukin-1beta-and tumor necrosis factor-alpha (TNFalpha)-dependent, and involves a TNFalpha receptor-associated factor-and NFkappaB-inducing kinase-dependent signaling mechanism. J Biol Chem. 2000;275(11):7918–24.

7. Patel NS, Paris D, Mathura V, Quadros AN, Crawford FC, Mullan MJ. Inflammatory cytokine levels correlate with amyloid load in transgenic mouse models of Alzheimer’s disease. J Neuroinflammation. 2005;2(1):9.

8. Mrak RE, Sheng JG, Griffin WS. Glial cytokines in Alzheimer’s disease: review and pathogenic implications. Hum Pathol. 1995;26(8):816–23.

9. Griffin WS, Stanley LC, Ling C, White L, MacLeod V, Perrot LJ, et al. Brain interleukin 1 and S-100 immunoreactivity are elevated in Down syndrome and Alzheimer disease. Proc Natl Acad Sci U S A. 1989;86(19):7611–5.

10. Blum-Degen D, Muller T, Kuhn W, Gerlach M, Przuntek H, Riederer P. Interleukin-1 beta and interleukin-6 are elevated in the cerebrospinal fluid of Alzheimer’s and de novo Parkinson’s disease patients. Neurosci Lett. 1995;202(1–2):17–20.

11. Cacabelos R, Franco-Maside A, Alvarez XA. Interleukin-1 in Alzheimer’s disease and multi-infarct dementia: neuropsychological correlations. Methods Find Exp Clin Pharmacol. 1991;13(10):703–8.

12. Yin Z, Raj D, Saiepour N, Van Dam D, Brouwer N, Holtman IR, et al. Immune hyperreactivity of Abeta plaque-associated microglia in Alzheimer’s disease. Neurobiol Aging. 2017;55:115–22.

13. Halle A, Hornung V, Petzold GC, Stewart CR, Monks BG, Reinheckel T, et al. The NALP3 inflammasome is involved in the innate immune response to amyloid-beta. Nat Immunol. 2008;9(8):857–65.

14. Taneo J, Adachi T, Yoshida A, Takayasu K, Takahara K, Inaba K. Amyloid beta oligomers induce interleukin-1beta production in primary microglia in a cathepsin B-and reactive oxygen species-dependent manner. Biochem Biophys Res Commun. 2015;458(3):561–7.

15. Guo H, Callaway JB, Ting JP. Inflammasomes: mechanism of action, role in disease, and therapeutics. Nat Med. 2015;21(7):677–87.

16. Olsen I, Singhrao SK. Inflammasome Involvement in Alzheimer’s Disease. J Alzheimers Dis. 2016;54(1):45–53.

17. Sheedy FJ, Grebe A, Rayner KJ, Kalantari P, Ramkhelawon B, Carpenter SB, et al. CD36 coordinates NLRP3 inflammasome activation by facilitating intracellular nucleation of soluble ligands into particulate ligands in sterile inflammation. Nat Immunol. 2013;14(8):812–20.

18. Heneka MT, Kummer MP, Stutz A, Delekate A, Schwartz S, Vieira-Saecker A, et al. NLRP3 is activated in Alzheimer’s disease and contributes to pathology in APP/PS1 mice. Nature. 2013;493(7434):674–8.

19. Coon KD, Myers AJ, Craig DW, Webster JA, Pearson JV, Lince DH, et al. A high-density whole-genome association study reveals that APOE is the major susceptibility gene for sporadic late-onset Alzheimer’s disease. J Clin Psychiatry. 2007;68(4):613–8.

20. Liu CC, Liu CC, Kanekiyo T, Xu H, Bu G. Apolipoprotein E and Alzheimer disease: risk, mechanisms and therapy. Nat Rev Neurol. 2013;9(2):106–18.

21. Wang JC, Kwon JM, Shah P, Morris JC, Goate A. Effect of APOE genotype and promoter polymorphism on risk of Alzheimer’s disease. Neurology. 2000;55(11):1644–9.

22. Holtzman DM, Herz J, Bu G. Apolipoprotein E and apolipoprotein E receptors: normal biology and roles in Alzheimer disease. Cold Spring Harb Perspect Med. 2012;2(3):a006312.

23. Rebeck GW. The role of APOE on lipid homeostasis and inflammation in normal brains. J Lipid Res. 2017;58(8):1493–9.

24. Huynh TV, Davis AA, Ulrich JD, Holtzman DM. Apolipoprotein E and Alzheimer’s disease: the influence of apolipoprotein E on amyloid-beta and other amyloidogenic proteins. J Lipid Res. 2017;58(5):824–36.

25. Fan YY, Cai QL, Gao ZY, Lin X, Huang Q, Tang W, et al. APOE epsilon4 allele elevates the expressions of inflammatory factors and promotes Alzheimer’s disease progression: A comparative study based on Han and She populations in the Wenzhou area. Brain Res Bull. 2017;132:39–43.

26. Gale SC, Gao L, Mikacenic C, Coyle SM, Rafaels N, Murray Dudenkov T, et al. APOepsilon4 is associated with enhanced in vivo innate immune responses in human subjects. J Allergy Clin Immunol. 2014;134(1):127–34.

27. Vitek MP, Brown CM, Colton CA. APOE genotype-specific differences in the innate immune response. Neurobiol Aging. 2009;30(9):1350–60.

28. Rodriguez GA, Tai LM, LaDu MJ, Rebeck GW. Human APOE4 increases microglia reactivity at Abeta plaques in a mouse model of Abeta deposition. J Neuroinflammation. 2014;11:111.

29. Mannix RC, Zhang J, Park J, Zhang X, Bilal K, Walker K, et al. Age-dependent effect of apolipoprotein E4 on functional outcome after controlled cortical impact in mice. J Cereb Blood Flow Metab. 2011;31(1):351–61.

30. Bennett RE, Esparza TJ, Lewis HA, Kim E, Mac Donald CL, Sullivan PM, et al. Human apolipoprotein E4 worsens acute axonal pathology but not amyloid-beta immunoreactivity after traumatic brain injury in 3×TG-AD mice. J Neuropathol Exp Neurol. 2013;72(5):396–403.

31. Tu JL, Zhao CB, Vollmer T, Coons S, Lin HJ, Marsh S, et al. APOE 4 polymorphism results in early cognitive deficits in an EAE model. Biochem Biophys Res Commun. 2009;384(4):466–70.

32. Pocivavsek A, Burns MP, Rebeck GW. Low-density lipoprotein receptors regulate microglial inflammation through c-Jun N-terminal kinase. Glia. 2009;57(4):444–53.

33. Colton CA, Brown CM, Cook D, Needham LK, Xu Q, Czapiga M, et al. APOE and the regulation of microglial nitric oxide production: a link between genetic risk and oxidative stress. Neurobiol Aging. 2002;23(5):777–85.

34. Huebbe P, Lodge JK, Rimbach G. Implications of apolipoprotein E genotype on inflammation and vitamin E status. Mol Nutr Food Res. 2010;54(5):623–30.

35. Guo L, LaDu MJ, Van Eldik LJ. A dual role for apolipoprotein e in neuroinflammation: anti- and pro-inflammatory activity. J Mol Neurosci. 2004;23(3):205–12.

36. Lynch JR, Tang W, Wang H, Vitek MP, Bennett ER, Sullivan PM, et al. APOE genotype and an ApoE-mimetic peptide modify the systemic and central nervous system inflammatory response. J Biol Chem. 2003;278(49):48529–33.

37. Shi Y, Yamada K, Liddelow SA, Smith ST, Zhao L, Luo W, et al. ApoE4 markedly exacerbates tau-mediated neurodegeneration in a mouse model of tauopathy. Nature. 2017;549(7673):523–7.

38. Brown MS, Goldstein JL. Suppression of 3-hydroxy-3-methylglutaryl coenzyme A reductase activity and inhibition of growth of human fibroblasts by 7-ketocholesterol. J Biol Chem. 1974;249(22):7306–14.

39. Kandutsch AA, Chen HW. Inhibition of sterol synthesis in cultured mouse cells by cholesterol derivatives oxygenated in the side chain. J Biol Chem. 1974;249(19):6057–61.

40. Brown MS, Goldstein JL. Cholesterol feedback: from Schoenheimer’s bottle to Scap’s MELADL. J Lipid Res. 2009;50 Suppl:S15–27.

41. Lund EG, Kerr TA, Sakai J, Li WP, Russell DW. cDNA cloning of mouse and human cholesterol 25-hydroxylases, polytopic membrane proteins that synthesize a potent oxysterol regulator of lipid metabolism. J Biol Chem. 1998;273(51):34316–27.

42. Russell DW. Oxysterol biosynthetic enzymes. Biochim Biophys Acta. 2000;1529(1–3):126–35.

43. Diczfalusy U, Olofsson KE, Carlsson AM, Gong M, Golenbock DT, Rooyackers O, et al. Marked upregulation of cholesterol 25-hydroxylase expression by lipopolysaccharide. J Lipid Res. 2009;50(11):2258–64.

44. Cyster JG, Dang EV, Reboldi A, Yi T. 25-Hydroxycholesterols in innate and adaptive immunity. Nat Rev Immunol. 2014;14(11):731–43.

45. Russell DW. The enzymes, regulation, and genetics of bile acid synthesis. Annu Rev Biochem. 2003;72:137–74.

46. Blalock EM, Buechel HM, Popovic J, Geddes JW, Landfield PW. Microarray analyses of laser-captured hippocampus reveal distinct gray and white matter signatures associated with incipient Alzheimer’s disease. J Chem Neuroanat. 2011;42(2):118–26.

47. Papassotiropoulos A, Lambert JC, Wavrant-De Vrieze F, Wollmer MA, von der Kammer H, Streffer JR, et al. Cholesterol 25-hydroxylase on chromosome 10q is a susceptibility gene for sporadic Alzheimer’s disease. Neurodegener Dis. 2005;2(5):233–41.

48. Morgan AR, Turic D, Jehu L, Hamilton G, Hollingworth P, Moskvina V, et al. Association studies of 23 positional/functional candidate genes on chromosome 10 in late-onset Alzheimer’s disease. Am J Med Genet B Neuropsychiatr Genet. 2007;144B(6):762–70.

49. Schjeide BM, McQueen MB, Mullin K, DiVito J, Hogan MF, Parkinson M, et al. Assessment of Alzheimer’s disease case-control associations using family-based methods. Neurogenetics. 2009;10(1):19–25.

50. Laumet G, Chouraki V, Grenier-Boley B, Legry V, Heath S, Zelenika D, et al. Systematic analysis of candidate genes for Alzheimer’s disease in a French, genome-wide association study. J Alzheimers Dis. 2010;20(4):1181–8.

51. Orre M, Kamphuis W, Osborn LM, Jansen AHP, Kooijman L, Bossers K, et al. Isolation of glia from Alzheimer’s mice reveals inflammation and dysfunction. Neurobiol Aging. 2014;35(12):2746–60.

52. Holtman IR, Raj DD, Miller JA, Schaafsma W, Yin Z, Brouwer N, et al. Induction of a common microglia gene expression signature by aging and neurodegenerative conditions: a co-expression meta-analysis. Acta Neuropathol Commun. 2015;3:31.

53. Matarin M, Salih DA, Yasvoina M, Cummings DM, Guelfi S, Liu W, et al. A genome-wide gene-expression analysis and database in transgenic mice during development of amyloid or tau pathology. Cell Rep. 2015;10(4):633–44.

54. Radde R, Bolmont T, Kaeser SA, Coomaraswamy J, Lindau D, Stoltze L, et al. Abeta42-driven cerebral amyloidosis in transgenic mice reveals early and robust pathology. EMBO Rep. 2006;7(9):940–6.

55. Yoshiyama Y, Higuchi M, Zhang B, Huang SM, Iwata N, Saido TC, et al. Synapse loss and microglial activation precede tangles in a P301S tauopathy mouse model. Neuron. 2007;53(3):337–51.

56. Knouff C, Hinsdale ME, Mezdour H, Altenburg MK, Watanabe M, Quarfordt SH, et al. Apo E structure determines VLDL clearance and atherosclerosis risk in mice. J Clin Invest. 1999;103(11):1579–86.

57. Sullivan PM, Mezdour H, Quarfordt SH, Maeda N. Type III hyperlipoproteinemia and spontaneous atherosclerosis in mice resulting from gene replacement of mouse Apoe with human Apoe*2. J Clin Invest. 1998;102(1):130–5.

58. Sullivan PM, Mezdour H, Aratani Y, Knouff C, Najib J, Reddick RL, et al. Targeted replacement of the mouse apolipoprotein E gene with the common human APOE3 allele enhances diet-induced hypercholesterolemia and atherosclerosis. J Biol Chem. 1997;272(29):17972–80.

59. Westover EJ, Covey DF. Synthesis of ent-25-hydroxycholesterol. Steroids. 2006;71(6):484–8.

60. Dick MS, Sborgi L, Ruhl S, Hiller S, Broz P. ASC filament formation serves as a signal amplification mechanism for inflammasomes. Nat Commun. 2016;7:11929.

61. Wannamaker W, Davies R, Namchuk M, Pollard J, Ford P, Ku G, et al. (S)-1-((S)-2-{[1-(4-amino-3-chloro-phenyl)-methanoyl]-amino}-3,3-dimethyl-butanoy l)-pyrrolidine-2-carboxylic acid ((2R,3S)-2-ethoxy-5-oxo-tetrahydro-furan-3-yl)-amide (VX-765), an orally available selective interleukin (IL)-converting enzyme/caspase-1 inhibitor, exhibits potent anti-inflammatory activities by inhibiting the release of IL-1beta and IL-18. J Pharmacol Exp Ther. 2007;321(2):509–16.

62. Goldstein JL, DeBose-Boyd RA, Brown MS. Protein sensors for membrane sterols. Cell. 2006;124(1):35–46.

63. Lembo D, Cagno V, Civra A, Poli G. Oxysterols: An emerging class of broad spectrum antiviral effectors. Mol Aspects Med. 2016;49:23–30.

64. Gold ES, Diercks AH, Podolsky I, Podyminogin RL, Askovich PS, Treuting PM, et al. 25-Hydroxycholesterol acts as an amplifier of inflammatory signaling. Proc Natl Acad Sci U S A. 2014;111(29):10666–71.

65. Chalmin F, Rochemont V, Lippens C, Clottu A, Sailer AW, Merkler D, et al. Oxysterols regulate encephalitogenic CD4(+) T cell trafficking during central nervous system autoimmunity. J Autoimmun. 2015;56:45–55.

66. Jang J, Park S, Jin Hur H, Cho HJ, Hwang I, Pyo Kang Y, et al. 25-hydroxycholesterol contributes to cerebral inflammation of X-linked adrenoleukodystrophy through activation of the NLRP3 inflammasome. Nat Commun. 2016;7:13129.

67. Pokharel SM, Shil NK, Gc JB, Colburn ZT, Tsai SY, Segovia JA, et al. Integrin activation by the lipid molecule 25-hydroxycholesterol induces a proinflammatory response. Nat Commun. 2019;10(1):1482.

68. Riemenschneider M, Mahmoodzadeh S, Eisele T, Klopp N, Schwarz S, Wagenpfeil S, et al. Association analysis of genes involved in cholesterol metabolism located within the linkage region on chromosome 10 and Alzheimer’s disease. Neurobiol Aging. 2004;25(10):1305–8.

69. Shibata N, Kawarai T, Lee JH, Lee HS, Shibata E, Sato C, et al. Association studies of cholesterol metabolism genes (CH25H, ABCA1 and CH24H) in Alzheimer’s disease. Neurosci Lett. 2006;391(3):142–6.

70. Guven G, Vurgun E, Bilgic B, Hanagasi H, Gurvit H, Ozer E, et al. Association between selected cholesterol-related gene polymorphisms and Alzheimer’s disease in a Turkish cohort. Mol Biol Rep. 2019;46(2):1701–7.

71. Shi Y, Holtzman DM. Interplay between innate immunity and Alzheimer disease: APOE and TREM2 in the spotlight. Nat Rev Immunol. 2018;18(12):759–72.

72. Zhu Y, Nwabuisi-Heath E, Dumanis SB, Tai LM, Yu C, Rebeck GW, et al. APOE genotype alters glial activation and loss of synaptic markers in mice. Glia. 2012;60(4):559–69.

73. Reboldi A, Dang EV, McDonald JG, Liang G, Russell DW, Cyster JG. Inflammation. 25-Hydroxycholesterol suppresses interleukin-1-driven inflammation downstream of type I interferon. Science. 2014;345(6197):679–84.

74. Dang EV, McDonald JG, Russell DW, Cyster JG. Oxysterol Restraint of Cholesterol Synthesis Prevents AIM2 Inflammasome Activation. Cell. 2017;171(5):1057–71 e11.

75. Bauman DR, Bitmansour AD, McDonald JG, Thompson BM, Liang G, Russell DW. 25-Hydroxycholesterol secreted by macrophages in response to Toll-like receptor activation suppresses immunoglobulin A production. Proc Natl Acad Sci U S A. 2009;106(39):16764–9.

76. Luo W, Liu W, Hu X, Hanna M, Caravaca A, Paul SM. Microglial internalization and degradation of pathological tau is enhanced by an anti-tau monoclonal antibody. Sci Rep. 2015;5:11161.

77. Molnar KS, Dunyak BM, Su B, Izrayelit Y, McGlasson-Naumann B, Hamilton PD, et al. Mechanism of Action of VP1-001 in cryAB(R120G)-Associated and Age-Related Cataracts. Invest Ophthalmol Vis Sci. 2019;60(10):3320–31.

78. Ramirez DM, Andersson S, Russell DW. Neuronal expression and subcellular localization of cholesterol 24-hydroxylase in the mouse brain. J Comp Neurol. 2008;507(5):1676–93.

