## Supplemental Figure 1 for "25-Hydroxycholesterol amplifies microglial IL-1β production in an apoE isoform-dependent manner"

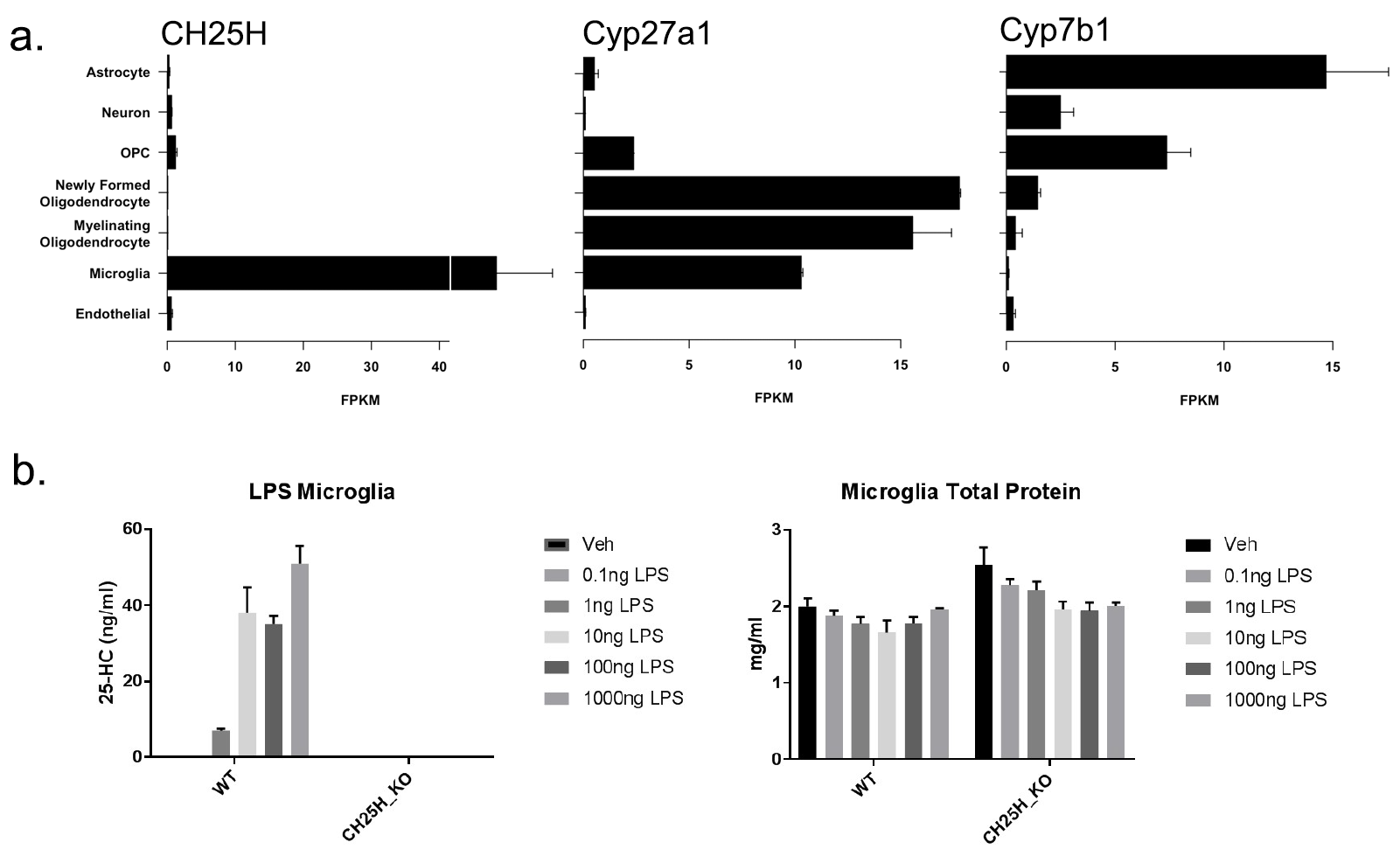

**Supplementary Figure 1.** a). Expression of CH25H, Cyp27a1 and Cyp7b1 in different cell types in brain based on the Stanford transcriptome database generated by Barres and colleagues (<http://www.brainrnaseq.org>). b). GC-MS analysis of 25-HC levels in the conditioned medium (left) and total protein levels of cell lysate of primary mouse microglia from wild-type and CH25H-/- mice treated with LPS (0, 0.1, 1, 10, 100, 1000ng/ml).
